# Estimating Visual Receptive Fields from EEG

**DOI:** 10.64898/2026.04.13.718144

**Authors:** Changxing Huang, Nanlin Shi, Yijun Wang, Xiaorong Gao

## Abstract

The visual receptive field (RF) characterizes the spatiotemporal properties of the visual pathway and serves as a fundamental unit for information encoding. While RFs have been extensively studied across various neural modalities, such as functional Magnetic Resonance Imaging (fMRI), Electrocorticography (ECoG), and Magnetoencephalography (MEG), their investigation via Electroencephalography (EEG) remains limited. In this study, we introduce a stimulation paradigm that combines white noise image sequences with a letter detection task to elicit central visual field EEG responses. Using the aligned/shuffled reverse correlation, we estimate RFs across different resolutions and demonstrate that the resulting RFs exhibit rich spatiotemporal characteristics. To validate the reliability of the estimated RFs, we constructed a visual EEG reconstruction model, which achieved good performance in a classification task. The same RF estimation method was subsequently applied to high-density EEG recordings to investigate the information gain afforded by high-density configurations in visual space. This work fills a gap in the study of visual RFs regarding the EEG modality and may inform the paradigm design of visual brain–computer interfaces.

## Introduction

The visual receptive field (RF), which reveals the spatiotemporal characteristics of the visual pathway^1^, has been widely studied by different modalities such as fMRI^2,3^, ECoG^4,5^, MEG^6–9^. From a single-neuron to the population level^10^, the RF may have variable temporal or spatial patterns^11,12^, and presents corresponding relationships with the anatomical structures of the cortex^13,14^. In addition to the aforementioned neural modalities, EEG also exhibits rich spatiotemporal information, which has been widely used in neuroscience and non-invasive brain-computer interface (BCI) research due to its ease of acquisition and high temporal resolution^15–17^. However, its relatively poor signal quality and low spatial resolution limit its application in more refined neural decoding studies^18^. This limitation is also evident in the field of visual receptive fields, where there are few reports directly estimating receptive fields using EEG^13,19,20^. This study employed a stimulation paradigm combining white noise image sequences with a letter detection task to elicit stimulus-locked visual EEG responses in the central visual field. Considering that the size of stimuli in the stimulus paradigm can affect the final

RF results^21^, we selected three different patch sizes of grids: 1°, 1.5°, and 2°. And then, we use aligned/shuffled reverse correlation^22^ to obtain the reliable spatio-temporal RF of different sensors, with various location and size. Compared with directly analyzing event-related potentials (ERPs) and visual evoked potentials (VEPs)^23–25^, this approach to deriving stable spatiotemporal patterns can reduce the impact of variability in EEG activity. This method has been demonstrated to effectively estimate receptive fields, with the resulting RFs exhibiting rich spatiotemporal characteristics.

The RF can also be understood as a response function of a time invariant system, which can be used for modeling neural pathways and predicting neural responses corresponding to input stimuli. Under white-noise stimulation, the time-domain result obtained via reverse correlation is equivalent to the system’s temporal response function (TRF)^26^. Recent studies have utilized RF models for visual image classification and reconstruction^27–29^, with some comparing RF models to deep learning models and attempting to interpret the mechanisms of layers^30,31^. In our previous work, we only considered the temporal characteristics of the RF and reconstructed EEG responses corresponding to a single white noise stimulus sequence^32,33^. Here, we perform dimensionality reduction on multi-channel EEG, extract the top four components to solve the spatiotemporal RF, and use the RF model corresponding to the first component to reconstruct and classify EEG activity under different white noise image sequences. Algorithms based on similar models may be used in the design of some BCIs based on spatial encoding paradigms in the future^34,35^.

This study aims both to fill the gap in EEG-based investigations of visual RFs and to offer theoretical insights for the design of visual BCIs. In addition, we seek to offer new perspectives on several ongoing research topics, such as high-density EEG. High-density EEG increases electrode density based on the original acquisition setup, aiming to enhance the spatial resolution of EEG signals^36^. It has demonstrated improved source localization performance^37–39^, and some studies on visual EEG have incorporated high-density modalities to enhance visual target classification performance^40^. In fact, high-density EEG recordings have been used to extract more detailed spatiotemporal information from visual responses^41,42^. However, few studies have directly illustrated the spatial information gain they provide in visual space. Therefore, we collected high-density EEG data under the same task, solved the reliable spatiotemporal RF of each channel, and compared the results of high-density (66 channels) and low-density (19 channels) from the perspectives of spatial coverage, gradient variance of spatial probability density, spatial information entropy, etc. Indeed, we observed improvements in several metrics with high-density EEG. In the future, high-density recordings could be applied to the decoding of complex visual scenes^43^ rather than single-object targets, enabling a more comprehensive exploration of the spatial information they offer—beyond merely enhancing the signal SNR.

## Results

Considering the characteristics of EEG signals, we employed white-noise image sequences combined with a letter detection task^5^ (Fig.1a) to elicit visual responses in the central visual field. The spatial contribution of visual input was characterized via stable temporal response patterns. Experiments were conducted using grids with three different patch sizes while maintaining comparable overall stimulus dimensions (Fig.1b). For each EEG channel, the original spatiotemporal receptive field (STRF) was estimated via reverse correlation with the corresponding stimulus sequence (20 sets, 6 repeats). In the time domain, this amounts to estimating a TRF for each electrode at each spatial location in the visual field. The spatial receptive field can then be characterized using the energy of these TRFs. However, noise and spontaneous activity in the EEG can cause false positive components in the visual space (see Supplementary Fig.1). To isolate stimulus-unrelated components, we performed reverse correlation after randomly shuffling the correspondence between EEG responses and stimuli. A spatial weight *W* was then derived by applying a threshold of 3 s.d. across multiple permutations (see Methods ‘*Aligned/Shuffled Reverse Correlation*’ for details), allowing us to obtain reliable STRFs (Fig.1c,d) and mitigate the impact of false positives on subsequent spatial analyses. Following prior studies^5,44^, we used the root-mean-square (RMS) map to represent the spatial RF, and parameters from a 2D Gaussian fit were used to quantify the RF’s center location and size (Fig.1e; see Methods ‘*RMS Map and Gaussian Fit’*). 15 participants completed the experiment using a standard EEG setup, while 5 participants (partially overlapping) participated in the high-density EEG condition. Fig.1f and Fig.1g present the STRFs (before weighting), RMS maps, and Gaussian fitting results for the O1 channel in the WN15 paradigm and the POz channel in the WN10 paradigm, respectively, from a representative subject. These results demonstrate that the proposed method can effectively extract the spatiotemporal features of RFs across different EEG channels while suppressing noise, yielding reliable estimates of RF location and size. Participants’ performance on the letter detection task across paradigms is shown in Fig.1h, confirming that all subjects maintained high attentional engagement.

**Fig. 1:**
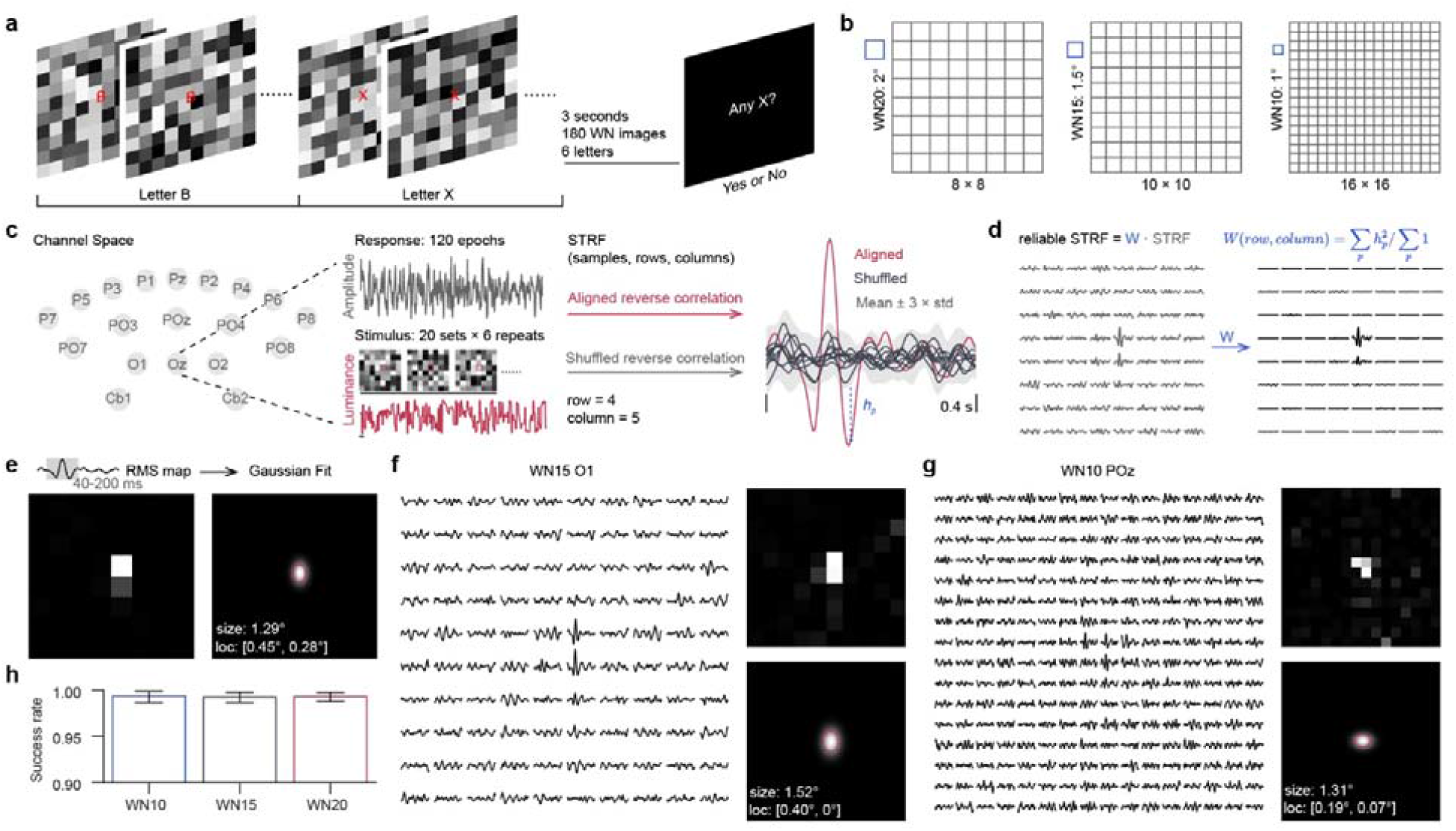
Experimental task and analysis pipeline. **a**, RF experiment combining white noise stimulation and letter detection task; 3-second trials with 180 images and 6 letters. Participants report whether the letter “X” appears at the end of each trial. **b**, Stimulus paradigms: WN20 (8×8 grid, 2° patches), WN15 (10×10 grid, 1.5° patches), WN10 (16×16 grid, 1° patches). **c**, RF estimation via aligned/shuffled reverse correlation from 120 epochs (20 stimulus groups × 6 repetitions) across 19 electrodes. The waveform on the right shows the TRFs of Oz at visual position row 4, column 5. **d**, Reliable RFs derived by applying spatial weights to original RFs. Each waveform spans 0.4 seconds and is normalized. **e**, RMS maps computed from the 40–200 ms interval are fitted with 2D Gaussians to estimate spatial position and size. The red ellipse indicates the FWHM contour of the fitted RF. **f**,**g**, Representative examples of original RFs, RMS maps, and Gaussian fits. Panel **f** shows the result from O1 under WN15; **g** shows the result from POz under WN10. **h**, Performance in the letter detection task. Bar plots represent the mean ± 95% CI; n=15.

### Characteristics of the estimated STRF

We first analyzed the characteristics of the estimated STRFs in 15 participants (Fig. 2, Supplementary Fig.2). To ensure the reliability of the spatial RF patterns, we excluded RFs with failed Gaussian fitting, as well as those located outside the central visual field (defined as within 8°) or had excessively large sizes (greater than 8°). Fig. 2a illustrates the spatial distribution of RFs across participants under different stimulation paradigms. The WN20 and WN15 paradigms retained more effective results (N = 209 for WN20; N = 179 for WN15; N = 285, with no exclusions), whereas only 122 remained under the WN10 paradigm. This suggests that the size of the stimulus content (patches) affects the reliability of the estimated RF. Large stimulus content can induce strong visual responses^40,45^, but may compromise spatial resolution of the visual field. The temporal and spectral patterns of the estimated RFs are illustrated in Fig.2b and 2c. The temporal dynamics were consistent with previous findings^32,33^, exhibiting prominent activity within the 0.04–0.2 s window, with slight variations across channels and subjects (see Supplementary Fig.3). In the frequency domain, the dominant components were primarily located within the 6–20 Hz range. Spatially, the distribution of RFs was predominantly confined to within a 4° diameter around the center of the visual field (Fig.2d). The proportion of RFs within this central area varied across paradigms: 82.3% in WN20, 89.4% in WN15, and 77.1% in WN10. In the context of visual BCI design^46,47^, the size of flickering stimuli is typically set to around 4°, and prior work has reported that stimulus-locked EEG response metrics under central fixation tend to approach a plateau when the central stimulus reaches ~3.8° (relative to peripheral stimulation)^45,48^. This qualitative pattern is consistent with the central concentration of RF locations observed in our study. Fig.2f illustrates the estimated RF sizes across different paradigms, with each dot representing the mean RF size across all effective results for an individual participant. Channel-wise results are provided in the Supplementary Fig.4. A clear positive correlation is observed between the estimated RF size and the patch size of the stimulus in each paradigm, and only a small fraction of the stimulus content appears to be captured at the channel level (WN20: 1.47° ± 0.31°, WN15: 1.21° ± 0.43°, WN10: 0.91° ± 0.39°). We interpret this as a consequence of the characteristics of EEG signals^49,50^, in which only components with strong responses or high SNRs are likely to be extracted. These components are typically associated with a limited subset of the stimulus elements.

**Fig. 2:**
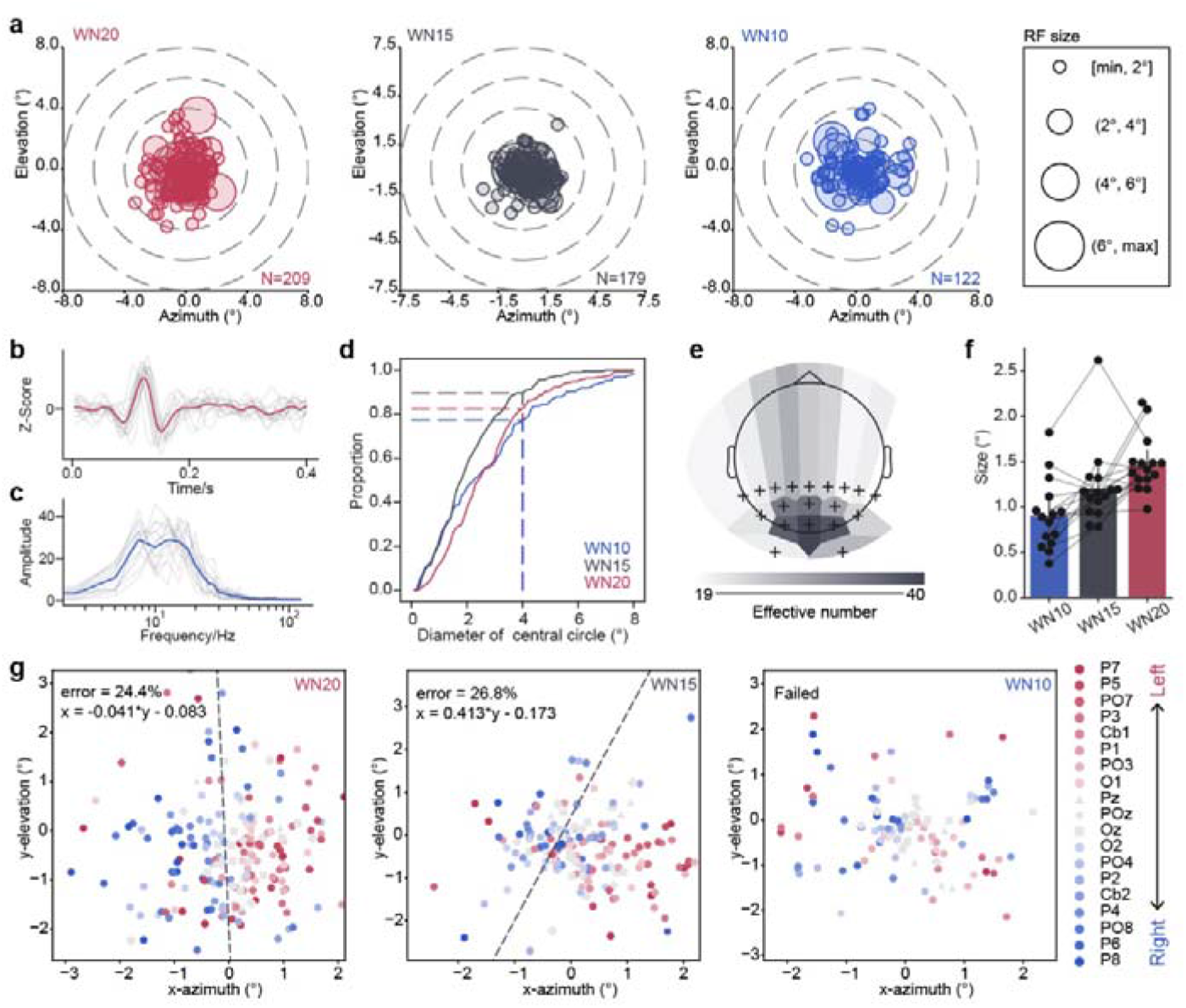
Characteristics of the estimated receptive fields. **a**, Spatial distribution of RFs across participants. Each circle represents the RF corresponding to one channel, with the radius proportional to the RF size as indicated by the legend on the right. N denotes the number of valid RFs under each stimulus paradigm. **b**, Time-domain RF responses from all participants at the location with the maximum visual spatial weight in the Oz electrode RFs. The gray lines represent individual participant waveforms; the red curve shows the group average. **c**, Frequency-domain RF responses corresponding to panel **b**. The blue curve indicates the average frequency response. **d**, Distribution proportion of RFs within circular regions centered on the visual field; the x-axis corresponds to the diameter of these circles. **e**, Topographical map of effective number for each channel across all paradigms, where ‘effective number’ denotes, for a given electrode, the total number of retained valid results (i.e., reliable RF estimates) aggregated across all participants. ‘+’ marks electrode positions; colors reflect the number of valid RFs per channel. **f**, RF size results across paradigms. Each point represents the average RF size across all valid RFs for one participant. Bars indicate mean ± 95% CI. **g**, Spatial distribution of RF locations across paradigms and LDA classification results. Each point shows the RF position in visual space. Left-hemisphere electrodes (odd-numbered names) are colored red; right-hemisphere electrodes (even-numbered names) are blue. The black dashed line indicates the LDA decision boundary.

### RF representations corresponding to cortical organization

In addition, we found that the estimated RFs exhibited a degree of correspondence with cortical organization. Fig.2e presents a topographic map of the number of effective RFs for each channel across the scalp. The highest counts were observed in occipital electrodes such as Oz, O1, O2, POz, PO3, and PO4, which typically capture the strongest stimulus-locked visual EEG responses. This finding suggests, on one hand, that these channels are indeed capable of capturing more robust visual EEG signals, and on the other hand, that our RF estimation approach can provide a coarse localization of the signal sources across the electrode array. Remarkably, the positions of EEG-based RFs revealed a retinotopically consistent mapping from the visual field to the cortex^51,52^. As shown in Fig.2g, we performed a linear discriminant analysis (LDA) to classify RFs derived from left-hemisphere channels (with odd-numbered labels) and distributed in the left visual field. The LDA decision boundary was defined as *x* = − 0.041*y* − 0.083, yielding a right-hemisphere channels (with even-numbered labels). In the WN20 paradigm, the RFs corresponding to left-hemisphere electrodes were predominantly located in the right visual field, while those from right-hemisphere electrodes were primarily classification error rate of 24.4%. In the WN15 paradigm, the LDA decision boundary shifted to *x* = 0.413*y* − 0.173, deviating from the ideal vertical midline (x = 0), with a slightly higher classification error rate of 26.8%. In the WN10 paradigm, the lateralized patterns were no longer distinguishable, and LDA failed to reliably separate the two groups. These results further indicate that the quality of RF estimation is closely related to the stimulus paradigm. Taken together, stimuli with patch sizes of 2° and 1.5° yielded more reliable visual RFs and better preserved the underlying retinotopic mapping between the visual field and cortical representations.

### Models after channel-space dimensionality reduction

In visual EEG studies, researchers commonly apply spatial filtering across electrode channels to enhance the SNR of the recorded signals^53–55^. Similarly, in our study, we employed Task Discriminant Component Analysis^56^ (TDCA; see Methods ‘*Spatial Filtering at the Channel Level*’) to perform spatial dimensionality reduction, extracting the top four components for RF estimation and subsequent reconstruction and classification tasks (Fig.3a). The spatial distributions of RFs in visual space under each paradigm are shown in Fig.3b. Following dimensionality reduction, the RFs associated with the principal components became more concentrated around the central visual field (see Supplementary Fig.5a), with the WN20 and WN15 paradigms retaining a greater number of effective results. To fully leverage the available data from each participant in the subsequent reconstruction and classification of visual EEG responses evoked by white noise stimuli, we examined the validity of the estimated RFs across components and paradigms. As shown in Fig.3c, all 15 participants yielded reliable RFs for the first component in both the WN20 and WN15 paradigms. Based on these results, we constructed RF models (Fig.4) for further analyses. Although the RFs derived from TDCA components exhibited more centralized spatial distributions compared to those based on original electrode channels, their estimated sizes remained largely consistent across paradigms (WN10: 0.97° ± 0.35°, WN15: 1.17° ± 0.44°, WN20: 1.46° ± 0.29°). Interestingly, we observed a positive correlation between the spatial weights of the TDCA filter (absolute values of the first component’s weights, indicating each channel’s contribution to the reduced signal) and the effective number of each channel (Fig.2e and Supplementary Fig.5b, c), suggesting that TDCA emphasized channels with stronger and more reliable visual responses. Fig.3e presents the topographic maps of TDCA weights under the WN20 and WN15 paradigms, showing that electrodes such as Oz and POz contributed the most to the first principal component. The relationship between TDCA weights and the effective numbers per channel is illustrated in Fig.3f. In both paradigms, a significant positive correlation was observed (WN20: *r* = 0.579, *p* = 9.37 × 10□^3^; WN15: *r* = 0.782, *p* = 7.80 × 10□□). This correlation can be interpreted from a SNR perspective. ‘Effective number’ is inherently linked to the reliability of the estimated RFs. In the process of estimating reliable RFs and computing the distance *h*_*p*_ to characterize spatial weighting across the visual field (see Methods), we assume that the signal comprises linearly explainable components. Under this assumption, responses to different stimuli are uncorrelated, and their means can be treated as a negligible constant. Then, this process can be simplified into an SNR structure (see Fig.3g and Supplementary Methods, where ***S***_*i*_ represents the signal components across channels in response to the *i*-th stimulus, 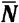 denotes the mean noise component, and ***Q***_*i*_ is the reverse correlation matrix for the *i*-th stimulus). Similarly, in the computation of TDCA weights across channels, we consider the trial-averaged responses as the signal component, and the residuals (trial-wise responses minus the average) as noise. This formulation can also be reduced to an SNR-like structure (see Fig.3g, where ***S***_*i*_ denotes the signal component for the *i*-th stimulus, and ***N*** is the noise matrix). The detailed derivation is provided in the Supplementary Methods. Previous studies have shown that these two definitions of the signal component correspond to the lower and upper bounds of the SNR^32,57^, respectively. The observed correlation between them suggests that the evoked visual EEG activity contains a substantial component that is linearly related to the stimulus.

**Fig. 3:**
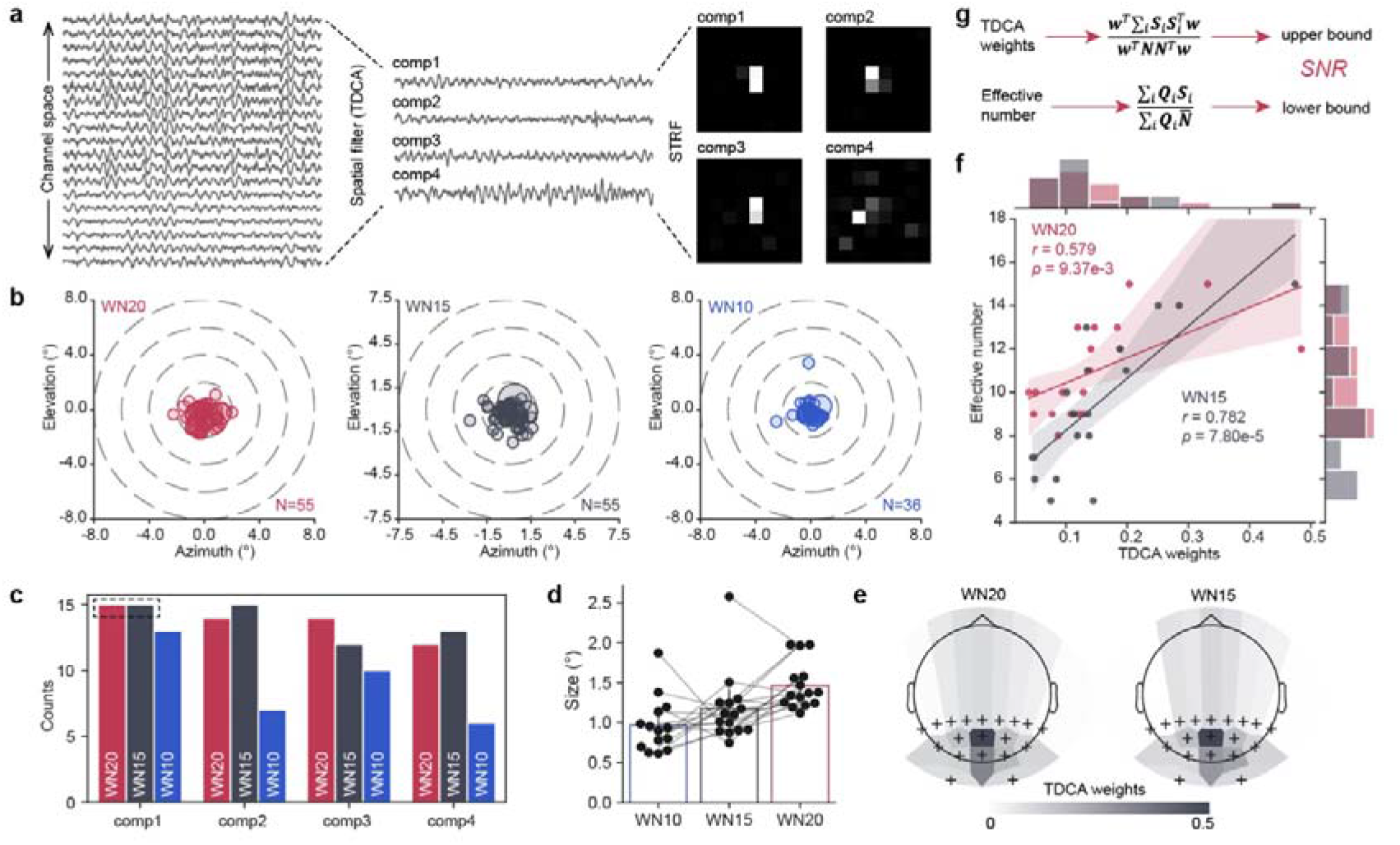
RFs after spatial filtering in channel space. **a**, Schematic of spatial filtering across channels using TDCA. The first four principal components are extracted, and RFs are estimated based on their spatial projections. **b**, Spatial distribution of RFs across participants. Each circle represents one RF corresponding to a principal component from one participant. **c**, Number of valid RFs across components and paradigms. The maximum value is 15 (number of participants). **d**, RF sizes across paradigms. Each dot represents the average RF size across all components for one participant. For one participant in the WN10 condition, no reliable RF estimates were obtained after spatial filtering. Bars indicate mean ± 95% CI. **e**, Topography of TDCA weights corresponding to the first component. ‘+’ symbols mark electrode locations; colors indicate weight magnitudes. **f**, Linear regression between effective number per channel and corresponding TDCA weights. Each dot represents one channel (n=15, error bar is 95% CI, WN20: *r*=0.579, *p*=9.37e-3; WN15: *r*=0.782, *p*=7.80e-5). **g**, Conceptual diagram illustrating the relationship between ‘effective number’, TDCA weights, and SNR.

### Visual response reconstruction and classification

To further evaluate the performance of the RF models, we conducted both reconstruction and classification of visual EEG responses elicited by different white noise image sequences. The data were first split into training and testing sets. Using the training data, we derived the TDCA spatial weights (first component) and the corresponding STRF. During testing, the stimulus sequences from the test set were convolved with the STRF to generate predicted responses (i.e., reconstruction templates; Fig.4b). The actual test EEG responses were then projected using the TDCA spatial filter, and their correlations with each template were computed. The label of the stimulus yielding the highest correlation was assigned as the predicted classification result (Fig.4a). To determine the minimum number of training data segments required to obtain a stable RF model, we evaluated the mean squared error (MSE) between the STRFs derived from the first *N* and *N* − 1 training segments (Fig.4c), with the data order randomly shuffled 20 times. We set a convergence threshold based on a pointwise difference of less than 0.1, corresponding to a total MSE below 0.01. Based on this criterion, a 10–10 split between training and testing sets was selected for the final reconstruction and classification analyses. Fig. 4d and 4e present the classification results under the WN20 and WN15 paradigms, respectively. We compared the performance of the original STRF (unweighted, see Fig.1d) with that of the reliable STRF, across different temporal window lengths. The results showed significant improvements when using the reliable STRF (*p* < 0.05 or *p* < 0.001, paired *t*-test), indicating that our approach indeed enhances the reliability of RF estimation. Among the top 20% of participants (ranked by classification performance), the accuracy achieved using the reliable STRF reached 91.1% ± 7.3% (WN20) and 83.8% ± 0.5% (WN15), whereas the original STRF yielded accuracies of 64.0% ± 18.6% (WN20) and 45.7% ± 8.5% (WN15). These findings demonstrate that EEG-based RF models can be effectively applied in visual BCI applications, and support the feasibility of more flexible and diverse paradigm designs. The correlation between the reconstructed and real EEG responses is shown in Fig.4f. A significant difference (*p* < 0.001, paired *t*-test) was observed between results obtained using the original STRF (light color) and the reliable STRF (dark color). Moreover, the correlations under the WN20 paradigm were significantly higher than those under the WN15 paradigm (F = 5.98, *p* = 0.0176, two-way ANOVA), consistent with the earlier receptive field analyses presented in Fig.2.

**Fig. 4:**
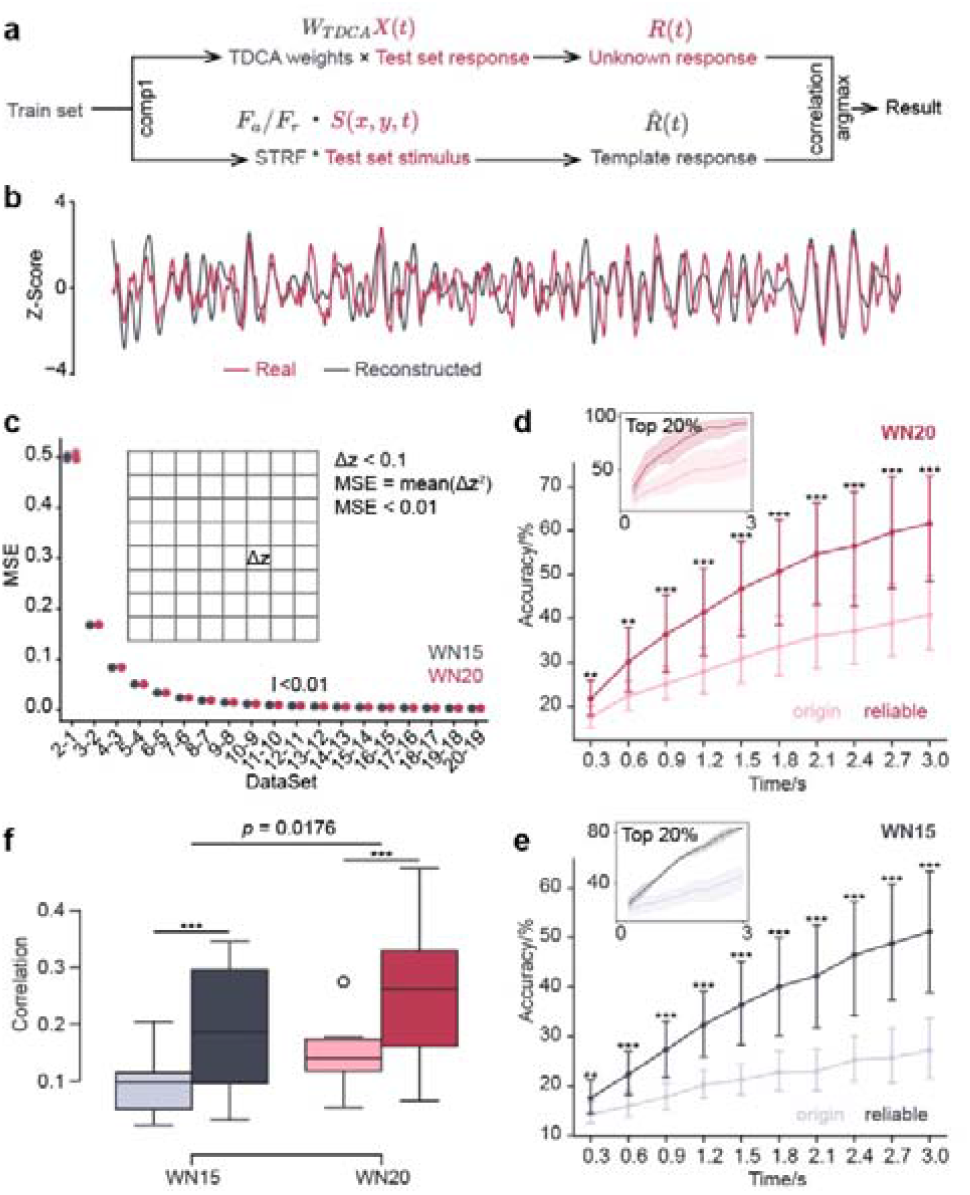
Reconstruction and classification of visual evoked EEG signals. **a**, Schematic of the visual EEG classification pipeline. TDCA weights and the STRF model (gray, from training data) are applied to testing data (red) for response classification. **b**, Example of EEG response reconstruction. The red curve shows the ground truth response; the gray curve represents the reconstructed response. Time window is 3 seconds. **c**, MSE between STRFs estimated from the first N and first N−1 stimulus groups. A threshold of MSE < 0.01 is used to determine the optimal split between training and testing data. **d**,**e**, Classification accuracy under the WN20 and WN15 paradigms, respectively. Each dot represents one participant. Insets show results from the top 20% performers. Bars indicate mean ± 95% CI (n = 15; paired t-test; \*\**p* < 0.01, \*\*\**p* < 0.001). **f**, Correlation between reconstructed and real responses for both paradigms (n = 15). Boxplots indicate medians, interquartile ranges, maxima, and minima. Light colors represent results from original STRFs; dark colors represent reliable STRFs. Statistical tests include paired t-test and two-way ANOVA; ***p < 0.001.

### Information gain from high-density recording

These findings motivated us to further apply the RF estimation approach to high-density EEG recordings, aiming to investigate whether increased spatial sampling yields additional visual field information. We conducted experiments on five participants using 66 occipital electrodes and extracted a subset of the original 19 electrodes for comparison. Under both the WN20 and WN15 paradigms, the resulting RMS maps for the 19-channel and 66-channel configurations are shown in Fig.5. The two configurations produced qualitatively similar spatial patterns within each paradigm, and the distribution of RF sizes showed no significant difference (Supplementary Fig.5d). While the 66-channel RMS maps appeared denser, they did not reveal substantial additional activation in other regions of the visual field. To quantitatively assess the impact of high-density recording, we modeled the distribution as a spatial probability density function *P*(*x,y*), and evaluated three spatial information metrics: spatial coverage, gradient variance, and entropy (see Methods ‘*Evaluation Metrics*’ for details).

**Fig. 5:**
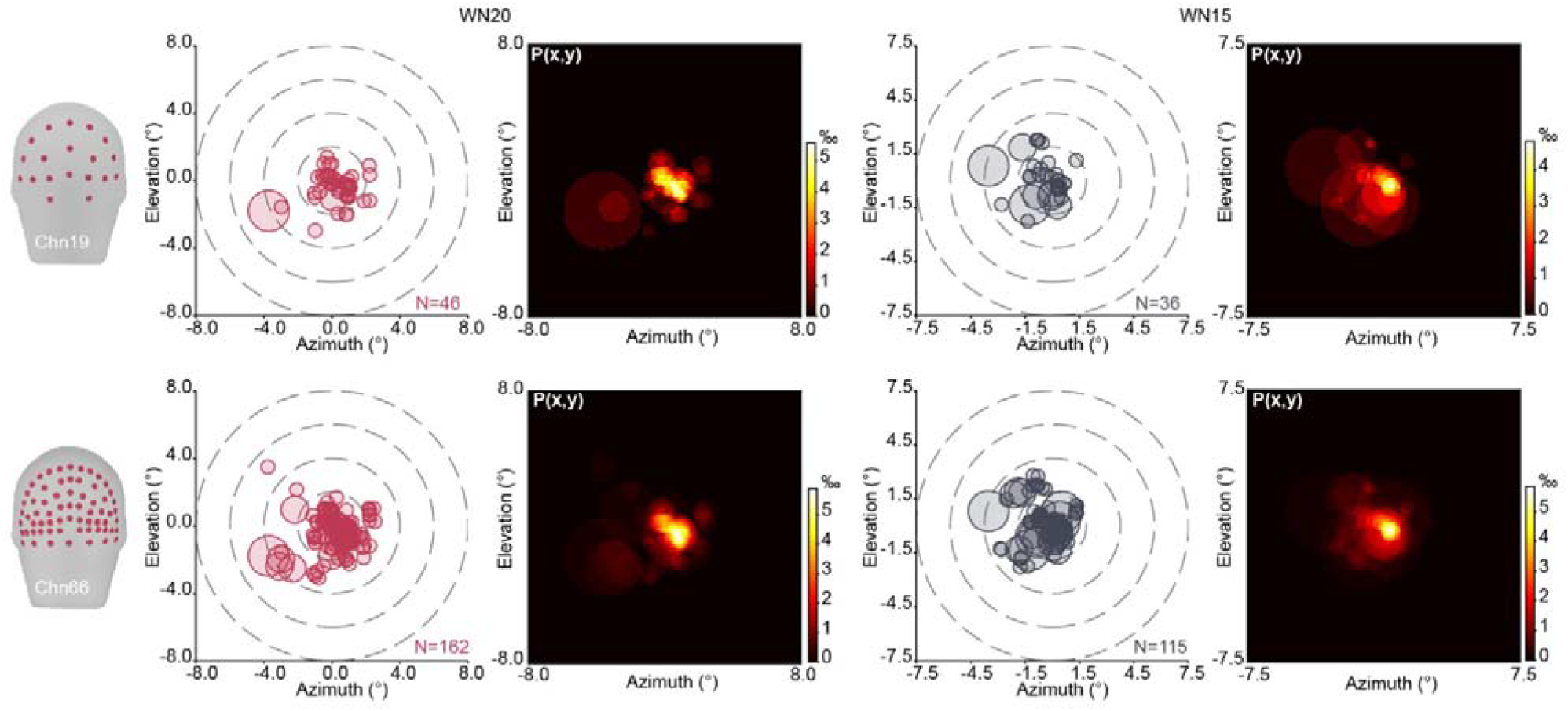
Spatial distribution of RFs under high-density EEG conditions. A matrix-style layout illustrates the spatial distribution of RFs and corresponding spatial probability density maps under two electrode configurations (Chn19 and Chn66) and two stimulus paradigms (WN20 and WN15). The RF maps follow the same visualization conventions as in previous figures.

Fig.6a illustrates the gain in spatial coverage provided by high-density EEG under both paradigms (WN20: 34.33% ± 5.25%; WN15: 50.32% ± 43.42%), suggesting that increased electrode density allows for a broader representation of the visual field. Additionally, the reduced gradient variance of the spatial probability density under high-density conditions (Fig.6b; WN20: –61.08% ± 20.81%; WN15: –60.24% ± 17.48%) indicates that high-density EEG yields smoother spatial representations in visual field, which may facilitate the decoding of finer visual details. To more comprehensively evaluate the spatial information gain afforded by high-density recordings, we calculated the information entropy of the spatial probability density (Fig.6c and 6d). Under the WN20 paradigm, high-density EEG showed a trend toward increased spatial entropy (*p* = 0.0625, Wilcoxon signed-rank test), while no significant difference was observed in the WN15 paradigm. We further performed reconstruction and classification of visual responses based on high-density EEG, using the same procedures as in Results ‘*Visual response reconstruction and classification*’ (employing reliable STRF models), and compared the results to those obtained from the 19-channel setup. In both paradigms, classification accuracy improved under the high-density condition, and the performance gap between the 66- and 19-channel configurations increased with longer temporal windows. However, due to the limited number of participants, these differences did not reach statistical significance (Fig.6f and 6g). A similar result was observed in the correlations between reconstructed and real responses (Fig.6e).

**Fig. 6:**
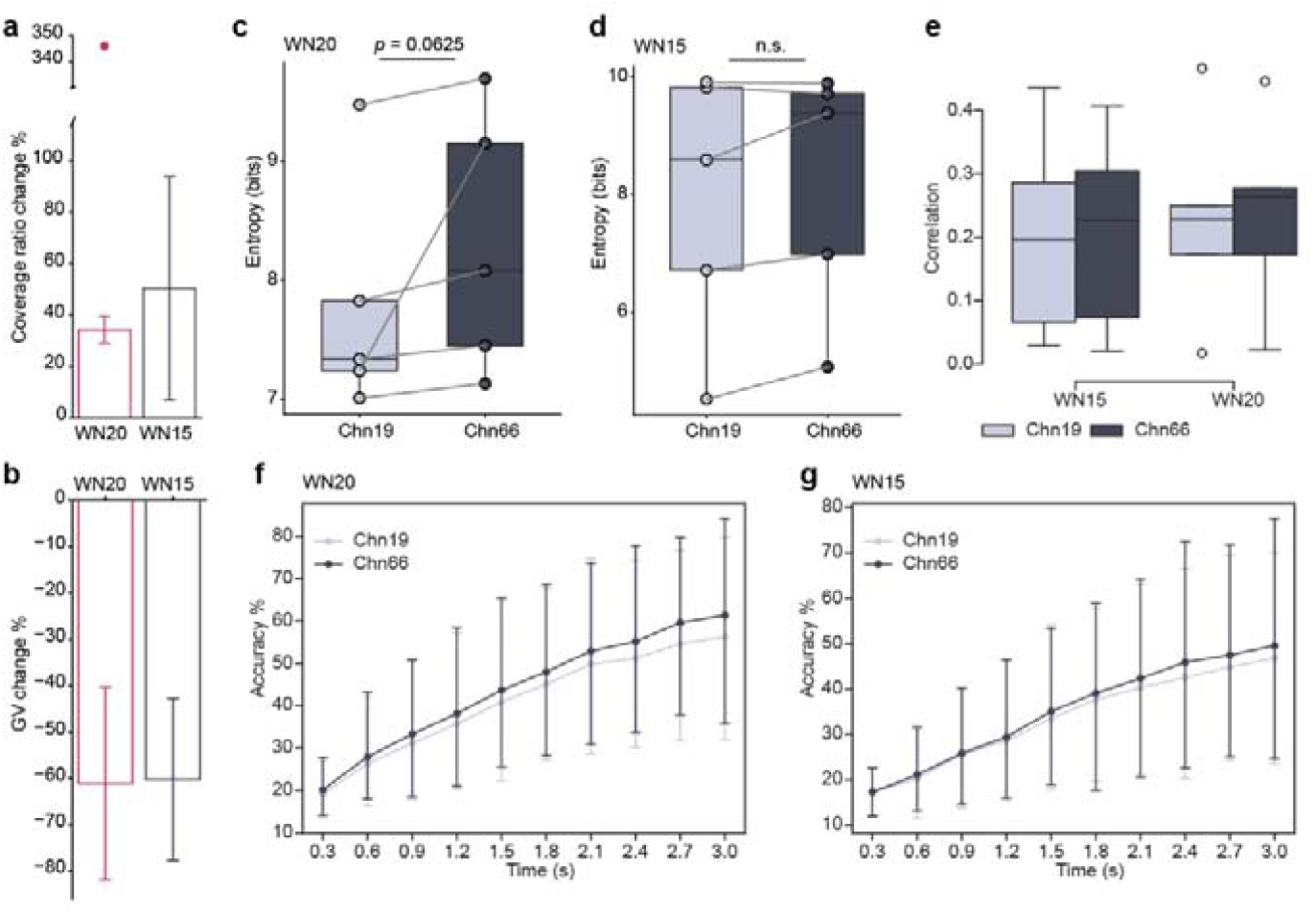
Information gain under high-density EEG configuration. **a**, Spatial coverage gain induced by high-density EEG across the two paradigms. **b**, Changes in gradient variance for both paradigms. In **a** and **b**, bars indicate mean ± std. Outliers are shown as individual points. **c**, Spatial information entropy under the WN20 paradigm. **d**, Spatial information entropy under the WN15 paradigm. In **c** and **d**, each dot represents one participant. Box plots show the interquartile range (Q1–Q3) with the median indicated; Wilcoxon signed-rank test is used for statistical analysis (n = 5, n.s. indicates no significance). **e**, Correlation between reconstructed and real responses (n = 5). Box plots display medians, interquartile ranges, maxima, and minima. **f, g**, Classification accuracy under the WN20 and WN15 paradigms, respectively. Error bars indicate 95% CI.

## Discussion

In this study, we established a method for estimating visual RFs under the EEG modality and applied it to both visual EEG reconstruction and high-density EEG analysis. We found that the spatial distribution of EEG-based RFs showed a coarse but meaningful correspondence with cortical organization, particularly in the lateralization between left and right visual fields. Unlike visual-field mapping approaches that rely directly on EEG VEP/ERP responses^23,25,58,59^, we estimate TRFs to capture the stable contribution of visual-field stimulation to the EEG response across space, thereby delineating, at the sensor level, the region of visual space to which each sensor is most responsive. In contrast to much of the prior work^58,59^, our primary goal is to construct a reliable STRF model that can support EEG encoding and decoding; consequently, the spatial constraints in our model may be more stringent. This is reflected in the estimated RF sizes. As discussed in the Supplementary Fig.6, although receptive fields are expected to become larger with eccentricity at the cortical level, a stimulus patch of fixed size does not engage an equally extensive cortical response in the peripheral field as it does near the center^58,60^. When reflected in scalp EEG, stimulus contributions under peripheral conditions are therefore weaker and less stable. As a result, the spatial visual-field extent that can be estimated reliably with our method tends to be smaller. Addressing this limitation may require leveraging deep-learning approaches^61,62^—for example, incorporating attention mechanisms—to capture richer information from the peripheral visual field. Nevertheless, we have established a relatively reliable STRF model that supports EEG reconstruction and classification, which contributes to visual EEG encoding/decoding, particularly in the context of visual BCIs. We found that the spatial distribution of RFs across channels was largely confined to within approximately 4° of the central visual field. After applying TDCA to extract principal components, this range was further reduced to around 3.5° (see Supplementary Fig.5a), which may also provide an intuitive explanation for how spatial filtering in multi-target speller BCI tasks^53–56^ can suppress the influence of surrounding stimuli. Moreover, our findings may inform the spatial layout of stimuli in visual BCI paradigms^63^ and our framework can provide algorithmic support for a broader range of spatially encoded visual BCI applications^34,64–67^. Finally, what advantages does high-density EEG offer over conventional EEG systems? In addition to its previously demonstrated benefits for BCI performance, our study showed that high-density EEG enhances spatial coverage, resolution, and entropy in receptive field representations. However, how these gains translate into improvements in real-world applications remains an open question that warrants further exploration.

To the best of our knowledge, no prior studies have directly estimated visual RFs from EEG. Since each EEG electrode captures aggregated neural activity and electrode positioning lacks high precision, the estimated RFs should be considered as coarse population RFs. Compared with fMRI^2,13^ and ECoG^5,44^, EEG-based RFs have lower physiological interpretability because they are derived from spatially pooled neural activity, and their correspondence to underlying cortical source components is less precise. The EEG-derived RF sizes are comparable in scale to the V1 RF sizes reported by Yoshor et al.^5^, which suggests that the stimulus-locked activity we capture may, to some extent, reflect contributions from early visual cortex. However, given the spatial pooling inherent to scalp EEG, contributions from other visual areas cannot be ruled out. These considerations highlight that how to more clearly characterize EEG receptive-field features associated with additional sources remains an important open challenge. We expect that neural network approaches may help address this issue. Despite these constraints, the noninvasive and efficient nature of EEG-based RF estimation offers considerable potential for clinical assessment of visual disorders and for noninvasive visual BCIs. Indeed, some recent clinical studies have attempted to detect visual field deficits from EEG responses^68^, underscoring the need for theoretical and technical advances in EEG-based RF estimation. We further suggest that EEG RFs may serve as a surrogate for ECoG in basic tasks. For instance, in the design of visual prostheses^69,70^, EEG RFs could be used for preliminary functional validation and coarse localization/stimulation within specific visual field regions, prior to subsequent ECoG implantation and experiments.

This study has several limitations. The RF estimation approach we employed is primarily linear. As shown in Supplementary Fig.7, the responses predicted by the linear model do not fully account for the observed EEG activity. In the future, incorporating nonlinear components^30^—either manually or through neural network-based approaches—may improve the system’s expressiveness and accuracy. Due to constraints of the stimulation design, using overly large stimulus grids reduces the spatial resolution of the estimated RFs in visual-field coordinates, whereas overly small grids fail to elicit sufficiently strong and stable responses. One possible refinement is to increase resolution by using overlapping stimuli^71^—for example, using 2° patches while shifting the patch centers by 1° between stimulus sets. Additionally, the reconstruction of visual responses in this study focused on EEG signals that had undergone spatial filtering to enhance the SNR. We did not attempt to reconstruct EEG responses at the level of individual channels. The current work primarily addresses the decoding process. However, once the STRF is obtained, the framework could, in principle, support not only EEG response reconstruction but also the inverse—reconstructing the stimulus image sequence from the EEG responses^72^. Realizing such a bidirectional mapping remains a challenging yet promising direction for future research.

## Methods

### Receptive Field Experiment and Analysis

#### Participants

A total of 15 participants (including 7 females) took part in the receptive field experiment under the standard electrode configuration. All participants had normal or corrected-to-normal vision and provided written informed consent prior to the experiment, in accordance with the protocol approved by the Institution Review Board of Tsinghua University. Participants received monetary compensation for their participation and agreed to the public sharing of anonymized EEG and behavioral data collected during the experiment.

#### Experimental Design

Participants were instructed to fixate on a central letter displayed on the screen, which changed every 0.5 seconds. At the end of each trial, they were asked to indicate whether the letter “X” had appeared. Meanwhile, the background was continuously updated at a frequency of 60 Hz using a sequence of white noise images, with each stimulus block was drawn from a uniform distribution between 0 and 1. Three types of visual stimulation resolution were employed: WN20 consisted of an 8 × 8 grid with each patch covering 2° of visual angle, WN15 used a 10 × 10 grid with 1.5° patches, and WN10 employed a 16 × 16 grid with 1° patches. The luminance of each patch was randomly sampled from a uniform distribution ranging from 0 to 1. For each paradigm, 20 unique sequences of 180 images (equivalent to 3 seconds of stimulation) were generated, and each sequence was repeated six times. Participants’ performance in the letter detection task was recorded and used to assess sustained attention throughout the experimental session.

#### Data Recording and Preprocessing

Visual stimuli were presented on an ASUS MG279Q monitor with a resolution of 1920 × 1080 pixels and a refresh rate of 60 Hz. The stimulus presentation was controlled using PsychoPy. EEG data were recorded using a Synamps2 system (NeuroScan Inc.) at a sampling rate of 1000 Hz, with signals acquired from 19 channels positioned according to the Quick-Cap Neo Net configuration, as shown in Fig.1c. The recorded EEG data were first downsampled to 250 Hz, followed by bandpass filtering between 4 and 100 Hz. Additionally, a notch filter was applied at 50 Hz to remove line noise interference.

#### Aligned/Shuffled Reverse Correlation

We first adopted the classical reverse correlation approach^22^ to estimate visual RFs in a linear framework. For each channel and each stimulus paradigm, we collected EEG responses (*R*) across 120 epochs, corresponding to 6 repetitions of 20 stimulus sequences (*S*). The original RF (*F*) was then computed according to the following equation:

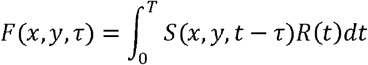

Here, *x* and *y* denote the spatial position of a stimulus patch within the white noise image, and *S* (*x,y,t*) represents the luminance value at location (*x,y*) and time point *t*. The total stimulus duration *T* for each trial was 3 seconds. The receptive field *F* was obtained by averaging across all trials. The temporal delay *τ* was defined over the range of 0 to 0.4 seconds^33^, allowing us to compute the aligned reverse correlation result *F*_*a*_ when the stimulus and EEG response were correspondingly paired. To mitigate the influence of spontaneous neural activity and random noise, we performed a group-wise temporal shuffling of the stimulus-response alignment, resulting in the shuffled reverse correlation output *F*_*s*_. This shuffling procedure was repeated 10 times. For each spatial position (*x,y*), the corresponding time series *f*_*s*_(*τ*) across iterations was used to compute the Mean_*s*_(*τ*) and SD_*s*_(*τ*). The aligned correlation response *f*_*a*_(*τ*)at each position was then compared against the shuffled distribution using the following equation:

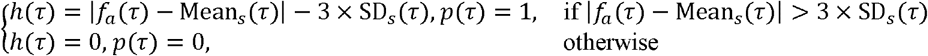

Based on these values, the spatial weight at each location (*x,y*)was defined as:

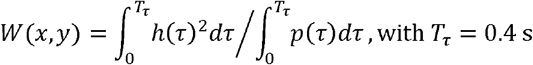

This weight reflects the average squared signal strength at a given spatial location, calculated only over time points (*p*) where the aligned results significantly exceeded the shuffled baseline (i.e., more than 3 standard deviations), *h* can be regarded as the distance from the TRF at *p* to the noise boundary (as shown in Fig.1c,d). The weights were then normalized across the entire visual field. The discrete version of this weighting procedure is illustrated in Fig.1c and 1d. Finally, the reliable receptive field *F*_*r*_ was computed by element-wise multiplying the aligned result *F*_*a*_ with the spatial weight matrix *w*:

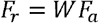

#### RMS Map and Gaussian Fit

To characterize the spatial properties of the RFs, we computed the RMS of each estimated RF within the 40–200 ms time window. Considering the robustness of the fitting results, we fitted a simplified two-dimensional Gaussian function (ignoring angular orientation) to each RMS map. The center coordinates of the fitted Gaussian were taken as the spatial position of the RF, while the average full width at half maximum (FWHM) along the two axes was used to define the spatial size of the RF^5,44^. Channels for which a valid Gaussian fit could not be obtained were labeled as ineffective, indicating that the signal from those channels did not contain sufficient information to support reliable receptive field estimation.

### Visual Response Reconstruction and Classification

#### Spatial Filtering at the Channel Level

We applied TDCA^56^ to perform spatial filtering across EEG channels. TDCA was used to extract principal component signals that capture discriminative features of the neural responses, and subsequent receptive field analyses were conducted based on these components. TDCA identifies a common spatial projection of EEG signals across channels by maximizing the between-class variance while minimizing the within-class variance. The objective of TDCA can be formally expressed as:

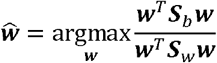

where ***w*** denotes the spatial weights across EEG channels (the absolute value of ***w*** is referred to as the TDCA weights), ***S***_*b*_is the between-class scatter matrix, and ***S***_*w*_ is the within-class scatter matrix. Detailed formulations of these matrices are provided in the Supplementary Materials.

#### Reconstruction and Classification Procedure

We selected the ***w*** corresponding to the first TDCA component as the spatial filter for downstream analysis. The EEG signal *X*(*t*) was projected through this filter to obtain the unknown response *R*(*t*) (see Fig.4a). The reconstruction response was generated by applying spatial summation and temporal convolution between the stimulus sequence and the STRF estimated from the first component.

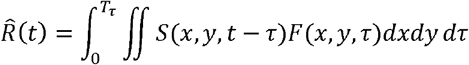

Here, *F* can be either *F*_*r*_ or *F*_*a*_, corresponding to the reliable or original RF, respectively (see Fig.4d). For each stimulus category (*i*),we computed a template response 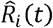 using the method described above. During testing, the unknown response *R*(*t*) was compared to all templates, and the category with the highest correlation coefficient was selected as the predicted label.

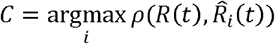

### High-Density EEG and Analysis

#### Experimental Setup

We used the Quick-cap Neo Net 256-channel configuration to record high-density EEG data and selected 66 electrodes located over the occipital region for RF estimation and analysis. The specific electrode selection^40^ is shown in the Supplementary Fig.8. A total of five participants (one female) took part in the high-density EEG experiment. Due to the lower reliability of RF estimation under the WN10 paradigm, only WN20 and WN15 paradigms were used in this part of the study.

#### Evaluation Metrics

To better characterize the spatial distribution of RFs in visual space under high-density conditions, we calculated the group-level spatial probability density *P*(*x,y*). Let *N*(*x,y*) denote the number of times the location (*x,y*) in visual space is covered by an individual receptive field. Here, *x* and *y* do not refer to the discrete positions of stimulus patches, but to a finer 100 × 100 point grid across the entire visual field. Then,

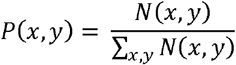

Furthermore, we introduced three metrics to compare the spatial information obtained under high-density and standard electrode configurations: spatial coverage ratio, gradient variance, and spatial entropy. Spatial coverage ratio was defined as the ratio between the area covered by RFs and the total visual field area. Gradient variance (*GV*) of the spatial probability density was used to evaluate the resolution of spatial information in the visual field. It was calculated as:

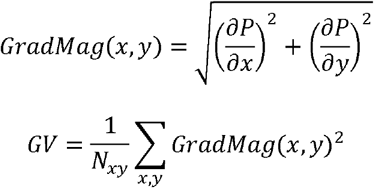

A smaller gradient variance indicates a smoother spatial distribution.

Spatial information entropy, as a global measure of spatial complexity, is defined as:

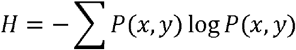

#### Statistical Analysis

All statistical analyses were performed using Python 3.9.13 (scipy 1.8.0, statsmodels 0.13.2). Specific statistical details are provided in the main text or figure captions. The methods used include paired t-tests, Wilcoxon rank-sum tests, and two-way ANOVA.

